# Beyond the Static Caliper: Dynamical Translocases and the Mathematical Imperative for Single-Molecule Proteomics

**DOI:** 10.64898/2026.07.20.739629

**Authors:** Jaylen E. Taylor, Parichit Sharma, Bryan A. Krantz

**Affiliations:** Department of Microbial Pathogenesis, School of Dentistry, University of Maryland, Baltimore, 650 W. Baltimore Street, Baltimore, MD 21201, U.S.A

**Keywords:** Single-molecule proteomics, nanopore sequencing, Physics-Informed Machine Learning, translocase, anthrax toxin, Markov kinetics, aerolysin

## Abstract

The advent of single-molecule nanopore sequencing established a powerful platform for modern genomics by using static biological pores to report the translocation of canonical nucleic acids, enabling rapid, accessible nucleic acid analysis. However, extending this strategy to single-molecule proteomics has stalled against a fundamental biophysical bottleneck. Current efforts in nanopore proteomics attempt to retrofit these static, spatial “caliper” biological nanopores (e.g., α-hemolysin, MspA, CsgG, aerolysin) for protein sequencing despite the immense steric, charge, and conformational heterogeneity of proteins. Unlike the chemically uniform, polyanionic phosphodiester backbone of DNA, the proteome contains isosteric and isobaric variants that confound purely volumetric measurements made by static pores. To address this bottleneck, we propose the application of *dynamical translocases* – naturally evolved, protein-handling nanomachines (e.g., the anthrax toxin protective antigen). Unlike static pores that rely on passive diffusion, dynamical translocases employ target-docking clamp architectures that achieve low nanomolar sensitivity. Active-site conformational dynamics generate high-dimensional kinetic fingerprints that enable molecular discrimination during translocation. By coupling dynamical translocases with Physics-Informed Machine Learning (PIML), we demonstrate that amino-acid side-chain-dependent thermodynamic friction can be mathematically decoded, enabling >90% accurate classification of chemically distinct amino acid classes and doing so label-free without the artificial DNA-handles required by legacy platforms.

## 1. The False Equivalence of Nucleic Acid and Protein Nanopore Sequencing

The success of single-molecule genomics emerged from a simple biophysical principle: passing a charged nucleic acid polymer through a rigid nanoscale aperture and recording the resulting electrical signal. Biological nanopores such as α-hemolysin ^1,2^, aerolysin ^3^, MspA ^4^, and CsgG ^5,6^ proved well suited as static “reading heads.” Because DNA relies on only four structurally distinct nucleobases appended to a predictable polyanionic phosphodiester backbone, the excluded volume of these bases within a rigid constriction is sufficient to generate discrete current blockade signatures. Buoyed by this success, the field has spent the last decade attempting to retrofit static DNA-sequencing pores for single-molecule proteomics. This effort, however, assumes that proteins can be interrogated using the same sensing architecture developed for nucleic acids. In contrast to DNA, the proteome is not a linear, uniformly charged polymer. Proteins comprise 20 canonical amino acids that are further diversified by post-translational modifications (PTMs), resulting in substantial variation in charge, hydrophobicity, and secondary structure. Attempting to resolve the chemical and conformational diversity of proteins using a static pore reduces the measurement to a single dimension: excluded volume within the rigid pore constriction. This fixed-geometry sensing paradigm is fundamentally constrained by the biological reality of the proteome. Static pores cannot distinguish between isosteric variants (e.g., stereoisomers of tryptophan) or isobaric residues (e.g., leucine and isoleucine), as they occupy an indistinguishable volume within the pore constriction. To compensate for these limitations, the field has increasingly relied on elaborate sample preparation strategies, including the covalent attachment of large polyanionic DNA handles that pull peptides through passive pores via helicase motors ^3,7^.

If single-molecule proteomics is to become a clinically deployable, label-free analytical platform, the field must move beyond the static DNA-sequencing paradigm. Resolving the chemical and conformational complexity of proteins, including their folded structures ^8^, requires a purpose-built ratcheting protein translocase engine ^9^. We propose that next-generation single- molecule proteomics relies on the deployment of *dynamical translocases* – complex, multi-component biological engines evolved specifically to capture, unfold, and report on complex polypeptide sequences with high sensitivity.

## 2. Dismantling the “Sensitivity Myth” (Active Capture vs. Passive Diffusion)

A common assumption within the single-molecule field is that smaller, rigid pores are inherently more “sensitive” than larger translocase complexes. This view conflates spatial constriction with target affinity, while overlooking the fundamental biophysics of molecular capture. To register a measurable translocation event, a target analyte must first locate and enter the pore vestibule. In established static platforms (e.g., MspA, CsgG, α-hemolysin), capture is dictated entirely by passive diffusion and stochastic collision. The probability of an unguided peptide navigating into a 1–2 nanometer aperture relies heavily on the bulk concentration of the analyte. Consequently, to achieve a statistically robust capture rate (events per minute) without the aid of processive DNA motor proteins, static pores often require analyte concentrations in the high micromolar (μM) to low millimolar (mM) range ^3,7^. In the context of clinical proteomics, where critical disease biomarkers often circulate at infinitesimal abundances, this requirement constitutes a critical bottleneck.

By contrast, biological translocases such as the anthrax toxin protective antigen (PA) ^10^ did not evolve to wait passively for stochastic collisions; they evolved to actively capture and transport low-abundance targets through an intricate, multi-tiered clamp architecture **(Fig 1A)**. The initial capture event is governed by the α-clamp ^11–15^, massive cleft structures that act as active docking stations. Rather than relying on simple diffusion, the α-clamp utilizes strong, non-covalent protein-protein interactions to actively capture and orient analytes from the bulk solution. Once docked, the analyte is sequentially guided into the ϕ-clamp ^13,15,16^ – a dynamic ring of phenylalanine residues – which ratchets the target through the transmembrane constriction. This active docking mechanism fundamentally rewrites the thermodynamics of capture. By converting a stochastic, diffusion-limited search into an affinity-driven docking event, the PA translocase drives the Limit of Detection (LOD) down into the low nanomolar (nM) to picomolar (pM) range ^16–27^. We and others have consistently demonstrated this ultra-sensitive capture of native peptides ^28^ and unfoldable domains ^29^. The notion that a large translocase complex is less sensitive than a passive DNA pore is inconsistent with the thermodynamics of molecular capture. The translocase is not a passive hole in a membrane; it is an active receptor, providing capture sensitivities orders of magnitude greater than those achievable with passive nanopores.

**Figure 1.**
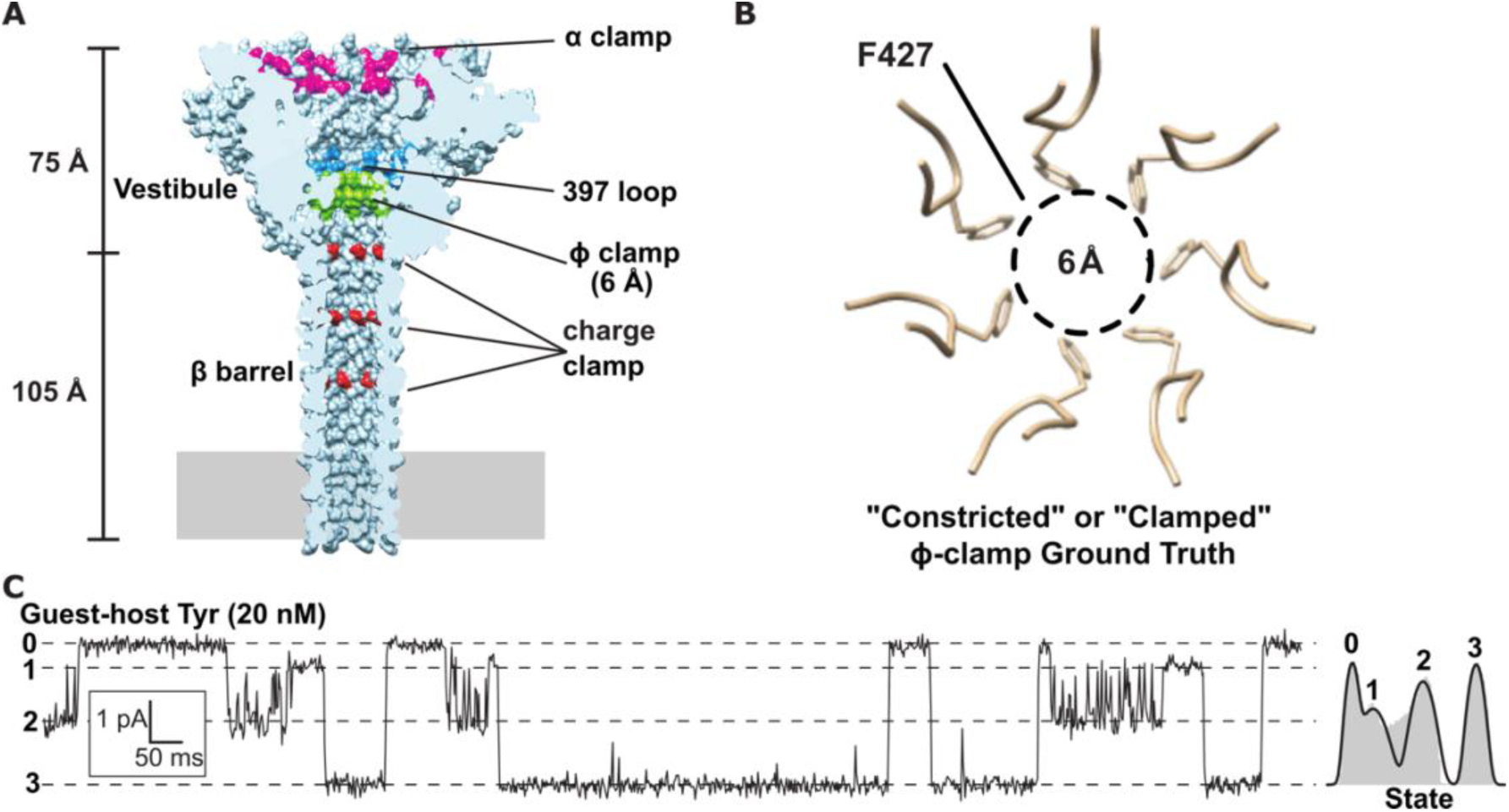
The Translocase Architecture: Redefining the “Reading Head.” **(A)** Sagittal cross-section of the PA nanopore derived from high-resolution Cryo-EM ^13,15^. The macroscopic ∼100 Å β-barrel and vestibule are denoted alongside numerous functional clamp and loop sites, illustrating the translocase’s structural capability to accommodate massive, folded polypeptides. **(B)** Inset highlighting the dynamic ϕ-clamp (Phe427) active site ^16^. The resting aperture in the cryo-EM structure exhibits an internal diameter of ∼6 Å, mathematically aligning with the van der Waals radius of a single amino acid side chain. This establishes that the primary electrical resistance and sensory capability of the channel are inherently localized to single-residue dimensions, definitively refuting the assumption that extended β-barrel architectures preclude high-resolution sensing. **(C)** Nanopore sensing guest-host calibration peptide standard: cis-added 20 nM guest-host Tyr peptide translocation events (sequence KKKKKYYSYY where guest = Y) at +70 mV (cis positive) in 100 mM KCl, pH 5.6. Discrete conductance states enumerated 0-3 are: fully blocked (State 0), partially ∼80-90% blocked intermediate 1 (State 1), partially ∼50% blocked intermediate 2 (State 2), and fully open (State 3). Logscale histogram on right is fitted to four Gaussians. Consistent data samples have been reported in Ghosal et al. ^21^, Colby and Krantz ^18^ and results described herein.

## 3. Dismantling the “Reading Head Fallacy” (Spatial Volume vs. Kinetic Transition)

The structural complexity of large translocase complexes has given rise to another common assumption: the “Reading Head Fallacy.” Critics often point to the overall dimensions of the PA heptamer or octamer, particularly its ∼100 Å-long transmembrane β-barrel and conclude that its structure precludes single-amino acid resolution. It is argued that a long barrel must encase multiple amino acids simultaneously, generating a blurred, unresolved signal. This criticism reflects a fundamental misunderstanding of the pore’s electrical circuitry, as the spatial length of the structural scaffold is decoupled from the dimensions of the electrical sensing region. In the PA translocase, the dominant electrical resistance – and therefore the primary source of the measurable ionic blockade – is highly localized to the ϕ-clamp (F427) at the top of the vestibule. Cryo-EM reconstructions ^13,15,30^ reveal that in its resting state, the ϕ-clamp possesses an internal diameter of approximately 6 Å **(Fig. 1B)**. This dimension is mathematically critical: it closely matches the van der Waals diameter of a single, bulky amino acid side chain. The peptide is not being indiscriminately “averaged” across a 100-Å barrel; the defining biophysical friction occurs precisely at this 6-Å bottleneck.

The “Reading Head Fallacy” relies on the assumption that the sensor is a rigid, spatial constriction. In contrast, the PA ϕ-clamp is a highly dynamic structure that actively ratchets and breathes – repeatedly clamping and unclamping peptide – during translocation ^20–23^, as is observed in single-channel electrophysiology recordings for all tested guest-host peptides **(Fig. 1C)** ^17–21^. As a peptide is ratcheted through the pore, the steric bulk of individual side chains physically forces the phenylalanine ring to dilate and constrict **(Fig. 2A)**. This dynamic interaction generates discrete structural micro-states (e.g., a fully blocked State 0, a deeply constricted State 1, and a dilated State 2) **(Fig. 1C)**. These breathing dynamics fundamentally change the nature of the measurement. We are not measuring the static excluded volume of a peptide in a pore; we are measuring the time-domain kinetic friction of a single amino acid actively perturbing a flexible gating mechanism, within a peptide- and nanopore-specific energy landscape **(Fig. 2B,C)**. These dynamics and energetics yield multi-dimensional kinetic fingerprints, characterized by highly specific dwell times, transition frequencies, and state probability ratios.

**Figure 2.**
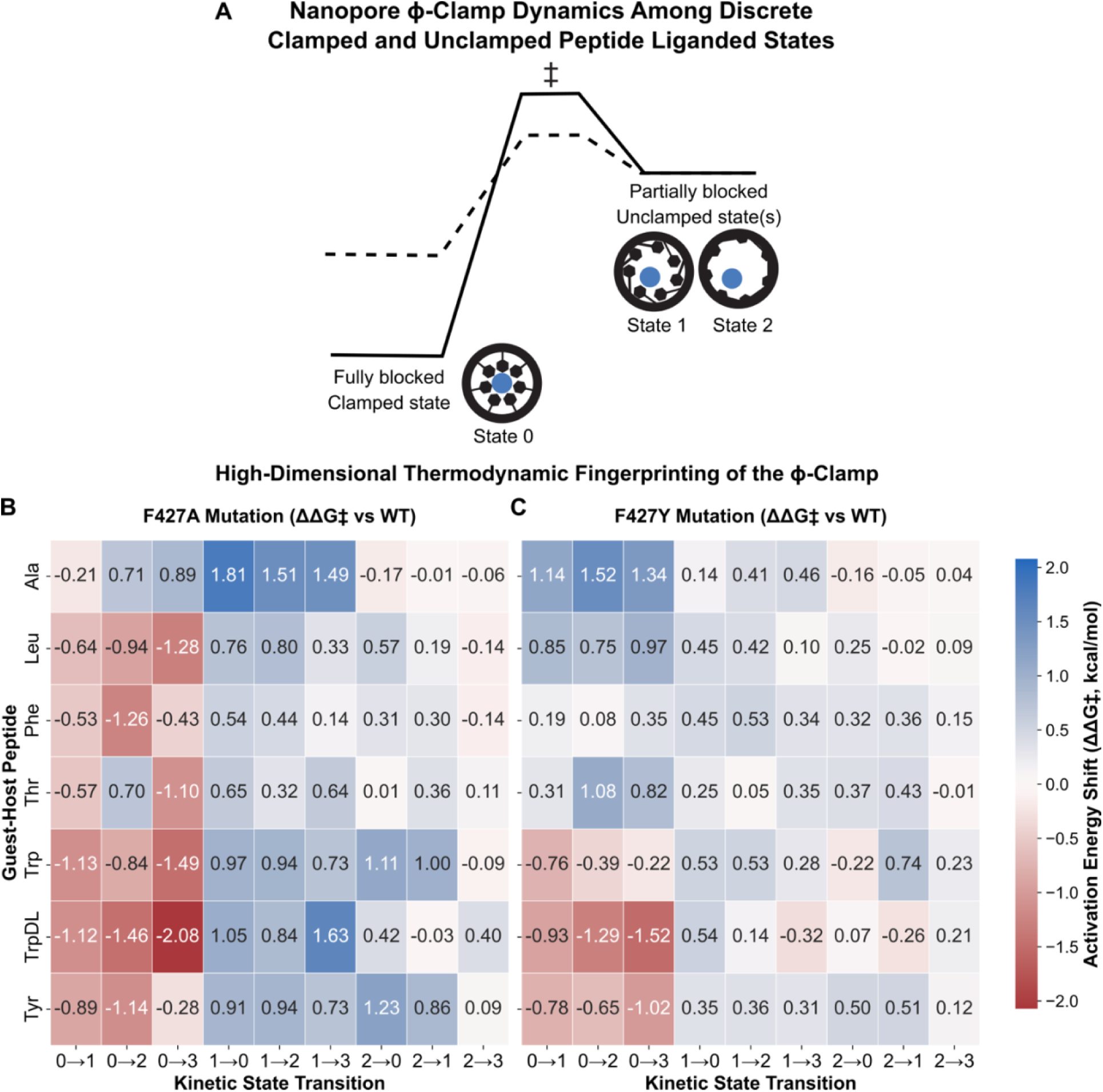
The High-Dimensional Thermodynamic Fingerprint of a Dynamical Sensor. **(A)** A conceptual energy landscape mapping the physical dilation of the PA ϕ-clamp to the time-domain kinetic signal. As a translocating peptide (blue circle) ratchets through the constriction, its steric bulk perturbs the F427 ϕ-clamp ring (black hexagons). The fully clamped, resting state corresponds to the highly resistive State 0, while structurally dilated conformations yield the partially blocked (∼80%, State 1) and unclamped (∼50%, State 2) intermediates. The dashed line illustrates the altered activation energy landscape of the F427A variant, demonstrating a modulated (shallower) energetic well at the reading head. **(B & C)** Quantitative matrices of activation energy shifts (ΔΔ*G*‡) for the **(B)** F427A and **(C)** F427Y ϕ-clamp variant translocases relative to wild-type PA across a select library of guest-host peptides. Rather than relying on 1D excluded volume, the dynamical translocase generates a high-dimensional thermodynamic signature. Red hues denote transition barriers that are energetically lowered (destabilized) by the mutation, while blue hues indicate increased (stabilized) transition barriers. The stark heterogeneity across the matrix mathematically demonstrates that every distinct amino acid sidechain perturbs the active site differently across multiple micro-state transitions. This complex, peptide-specific multi-dimensional friction provides the rigorous physical foundation required for PIML classification, a feat impossible in rigid, static-hole architectures. Data derived from the kinetic analyses of Colby and Krantz ^20^.

The PA translocase possesses a resolving power that surpasses that of static systems because it relies on kinetic transitions rather than pure spatial displacement. A rigid pore, for example, would not be able to differentiate between ʟ-Trp and ᴅ,ʟ-Trp peptides, as their spatial volumes are identical. We have empirically demonstrated, however, that the dynamic PA ϕ-clamp robustly resolves these stereoisomers ^17,18,21^, as their distinct topologies impose different thermodynamic friction on the active site, resulting in characteristic changes in gating frequency and kinetic state occupancy. Secondary-structure-dependent interactions may further contribute to this discrimination ^22,23^. Similarly, we achieve robust discrimination of isobaric residues (leucine vs. isoleucine), a challenge even for mass spectrometry, using a baseline machine learning classifier model (see below). The overall size of the PA translocase is not a hindrance to resolution; the dynamic nature of its internal gating mechanism is the prerequisite for achieving sub-Angstrom proteomic discrimination.

## 4. Dismantling the Poly-charged Leader Sequence Requirement (The “Lysine Trap”)

The development of single-molecule sensing has historically required highly charged, synthetic calibration standards ^17–21,26^. In the context of dynamical translocases, preliminary calibration often utilizes “guest-host” peptide models featuring a poly-cationic (e.g., poly-lysine) leader sequence. This design specifically isolates the kinetic signal of a target “guest” residue by providing an extremely rapid, highly uniform electrophoretic pull (Δψ) through the pore. Unfortunately, this foundational biophysical calibration strategy is often conflated with a biological limitation, giving rise to the pervasive “Lysine Trap” critique.

The notion that translocases such as PA require large polycationic leader sequences for translocation has contributed to the perception that they are incapable of reading diverse clinical biomarkers. This interpretation is inconsistent with both the biological design of the PA channel and the established physics of nanopore Δψ/ΔpH driving forces. As a dynamical nanopore, PA naturally unfolds and transports massive proteins of diverse sequence compositions, including large heterologous substrates approaching ∼90 kDa enzymes, across model planar bilayers or into host cells ^16,25,29,31–35^. To enable biologically native peptides to enter and transit a passive, rigid pores such as MspA or CsgG, the field has increasingly adopted elaborate sample-preparation strategies. Because static pores lack an intrinsic target affinity, peptide translocation often requires large polyanionic DNA handles to orient peptides within the electrical field or rely on processive DNA helicase motors to forcibly ratchet the target molecule through the constriction ^7,36^.

By contrast, the PA translocase operates free from these artificial constraints. Evolution did not design the PA channel to transport poly-lysine tracts; it designed it to unfold and transport, ∼90 kDa native virulence factors (i.e., Lethal and Edema Factors). Crucially, Lethal Factor is a net-anionic protein with an acidic isoelectric point (pI ∼5.5); as pointed out previously, the translocation of this large anionic enzyme via the cation-selective PA nanopore was a paradox entirely solved by differential protonation biophysics ^31^. Moreover, the PA translocase effortlessly processes this massive anionic cargo in seconds, driven entirely by the mildly acidic physiological proton gradient (ΔpH) of the host endosome ^31,32^. To fully appreciate the functional capacity of the PA translocase, one must examine the evolutionary bioinformatics of its native cargo. Information-theoretic analysis reveals that these virulence factors possess a biased amino-acid composition relative to the broader proteome. Whereas the global proteome exhibits a Shannon entropy of ∼4.18 bits per residue, mature LF and EF display substantially lower entropic states (4.01 and 3.98 bits, respectively), consistent with evolutionary adaptation for efficient translocation through the PA channel (**Fig. 3**).

**Figure 3.**
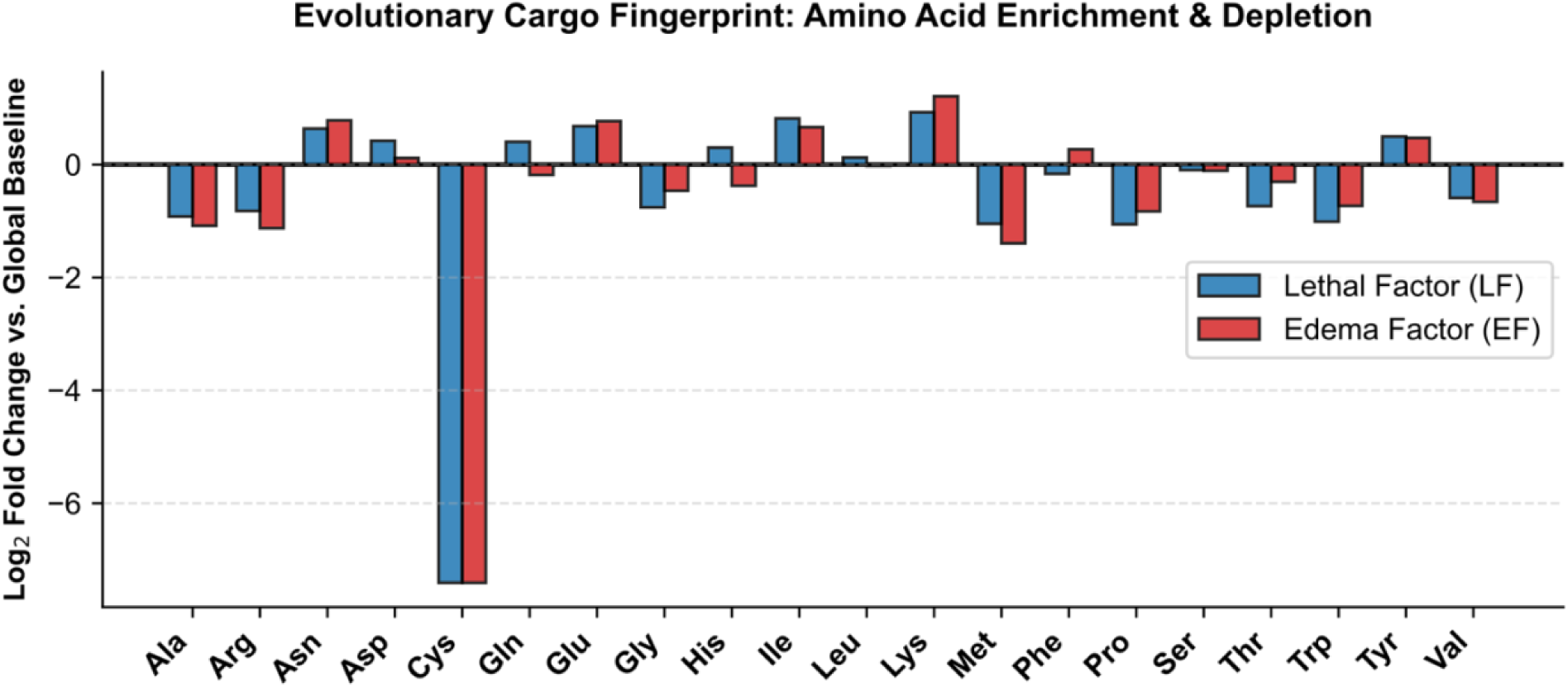
The Evolutionary Fingerprint of Translocase Cargo. Log_2_ fold-change of amino acid frequencies in mature *Bacillus anthracis* Lethal Factor (LF) and Edema Factor (EF) relative to the global proteomic baseline ^40^. Information-theoretic analysis reveals that mature LF (776 residues) and EF (767 residues) possess significantly lower Shannon entropy (4.01 and 3.98 bits, respectively) compared to the global baseline (4.18 bits). This compositional bias represents a targeted evolutionary optimization for nanopore translocation. Cysteine is entirely depleted to preclude pore-jamming disulfide bond formation. Tryptophan is heavily suppressed to avoid massive conformational entropy penalties within the narrow ϕ-clamp. Furthermore, Glutamate (higher pK_a_) is vastly enriched over Aspartate (lower pK_a_); this differential protonation capacity allows the cargo to more readily neutralize its charge in the acidic endosome, minimizing electrostatic friction during transit. This fingerprint demonstrates that the PA translocase is a highly tuned thermodynamic engine designed to process structurally “lubricated” biopolymers, contrasting sharply with passive, static nanopores.

This evolutionary signature is reflected in the selective enrichment and depletion of amino acid that influences energetic barriers to translocation. Cysteine is strongly depleted from the mature cargo to preclude covalent disulfide knots that would irrevocably jam the pore. Tryptophan, a bulky, rigid indole ring that incurs a significant conformational entropy penalty upon entering the 6 Å ϕ-clamp, is severely suppressed to prevent thermodynamic stalling. Conversely, Glutamate is vastly enriched over its structural counterpart, Aspartate. This bias is rooted in the thermodynamics of differential protonation. Native translocation is driven by the acidic gradient (ΔpH) of the host endosome ^31^. Because the sidechain pK_a_ of Glutamate (4.3–4.5) is notably higher than that of Aspartate (3.9–4.0), Glutamate is more readily protonated—and thus electrostatically neutralized—within the acidic micro-environment of the channel. This bias is consistent with evolutionary adaptation toward a polymer capable of shedding its negative charge, minimizing electrostatic friction against the translocase to facilitate rapid, fluid transit. Attempting to sequence native, unmodified proteins using static pores neglects the evolutionary adaptations that have optimized PA-mediated protein translocation over billions of years.

There is no structural or biological limitation preventing the PA translocase from processing anionic, neutral, or mixed-charge targets. The requirement is simply thermodynamic: translocation requires a driving force. For mixed-charge or cationic standard “bottom-up” proteolytic fragments, a standard electrophoretic field (Δψ) provides robust, label-free capture and transport. For strictly anionic or neutral fragments, the PA translocase seamlessly reverts to its native biological driving force (ΔpH). Synthetic DNA handles or helicase motors are not required to process relevant sequences using PA nanopores. The use of a poly-cationic leader is a deliberate, mathematically precise tool for single-channel calibration; interpreting it as a fundamental limitation of the translocase architecture misrepresents the underlying biophysics of the system.

## 5. The Hardware Bandwidth Imperative

The reliance on static pores and purely volumetric sensing architectures introduces a severe and often overlooked hardware bottleneck: the bandwidth. Because passive pores rely on minute variations in static excluded volume (*I_b_/I_0_*) to distinguish analytes, effective discrimination must sample electrical traces at extreme frequencies—often between 250 kHz and 1 MHz—to mathematically suppress the thermal noise floor. While achievable on isolated, single-channel patch-clamp measurements, such multi-megahertz sampling is difficult to scale to highly parallel CMOS architectures required for clinical proteomics. A commercial CMOS array containing 10,000 to 1,000,000 sensors would face substantial data-transfer, storage, and processing challenges at multi-megahertz sampling rates.

Conversely, dynamical translocases unburden the electrical hardware by shifting the discriminative work to the biological machine. Because the PA ϕ-clamp utilizes the thermodynamic friction of the peptide to generate discrete, analyte-dependent conformational state transitions, the critical biophysical telemetry – Markovian transition matrices and dwell times – manifests on the millisecond timescale. We have empirically demonstrated >90% classification accuracy across a seven-peptide set using data downsampled to 400 Hz ^17,18^. By requiring fractionally lower data bandwidths, dynamical translocases provide a mathematically viable route to scaling high-throughput, multiplexed nanopore arrays for commercial proteomics.

## 6. Quantitative Benchmarking: The Mathematical Ceiling of the Static-Pore Sensing

The true performance limits of static-pore sensing are often obscured by the lack of rigorous, unsupervised benchmarking. Claims of high-fidelity amino-acid recognition in static biological pores such as aerolysin are frequently supported by manually curated 1D amplitude histograms rather than automated, unsupervised classification frameworks ^3^. However, these visualizations can obscure the substantial pre-processing required to extract discriminative information from passive, volumetric measurements. To establish an objective empirical benchmark, we subjected published high-bandwidth (250 kHz) 20-amino-acid aerolysin datasets ^3^ to an automated, unsupervised Physics-Informed Machine Learning (XGBoost) pipeline. The targeted analyte was an XR_7_ guest-host peptide construct in which X represented each of the 20 canonical amino acids.

Analysis of the complete, uncurated dataset revealed substantial capture heterogeneity that is often obscured by curated histogram-based representations, what we term the “Data Curation Fallacy.” Without an active docking mechanism such as the α-clamp of PA, static pores rely entirely on passive stochastic diffusion, resulting in highly unequal capture rates across analytes. In equivalent recording windows, the dataset yielded over 20,000 arginine events but fewer than 250 tryptophan events. Furthermore, the uncurated event streams contained numerous sub-millisecond “incomplete collisions” and anomalous, multi-second intrinsic gating traps that required manual filtering prior to construction of 1D amplitude distributions. When our XGBoost meta-classifier was trained on the complete uncurated dataset (>56,000 events) and weighted to correct for the class imbalance, the overall 20-class accuracy reached 62.38%. To evaluate the influence of event curation on classification performance, we applied a strict >0.5 ms minimum dwell-time filter, mimicking manual preprocessing by excluding fast collisions to provide the classifier with the cleanest possible physical signal. Following application of the >0.5 ms dwell-time filer, the overall accuracy improved only modestly, increasing from 62.38% to 66.92% **(Fig. 4)**. The limited improvement achieved through increasingly aggressive preprocessing suggests that classification performance is constrained by the underlying physics of the measurement rather than by the data density alone. Because static pores cannot dynamically “breathe” through analyte dependent multistate transitions, the classifier lacks the high-dimensional kinetic transition features necessary to distinguish between isosteric and isobaric residues. Consistent with the limited discriminatory information available in the measurement, several amino acids remained poorly resolved with F1-scores of 0.12 for methionine, 0.16 for leucine, and 0.20 for glutamate. Their physical signals sit directly on top of each other in 1D excluded -volume space, providing limited discriminatory information for machine-learning classifiers. This empirical benchmarking suggests that true *de novo* sequencing is unlikely to be achieved through label-free, passive, volumetric measurement alone.

**Figure 4.**
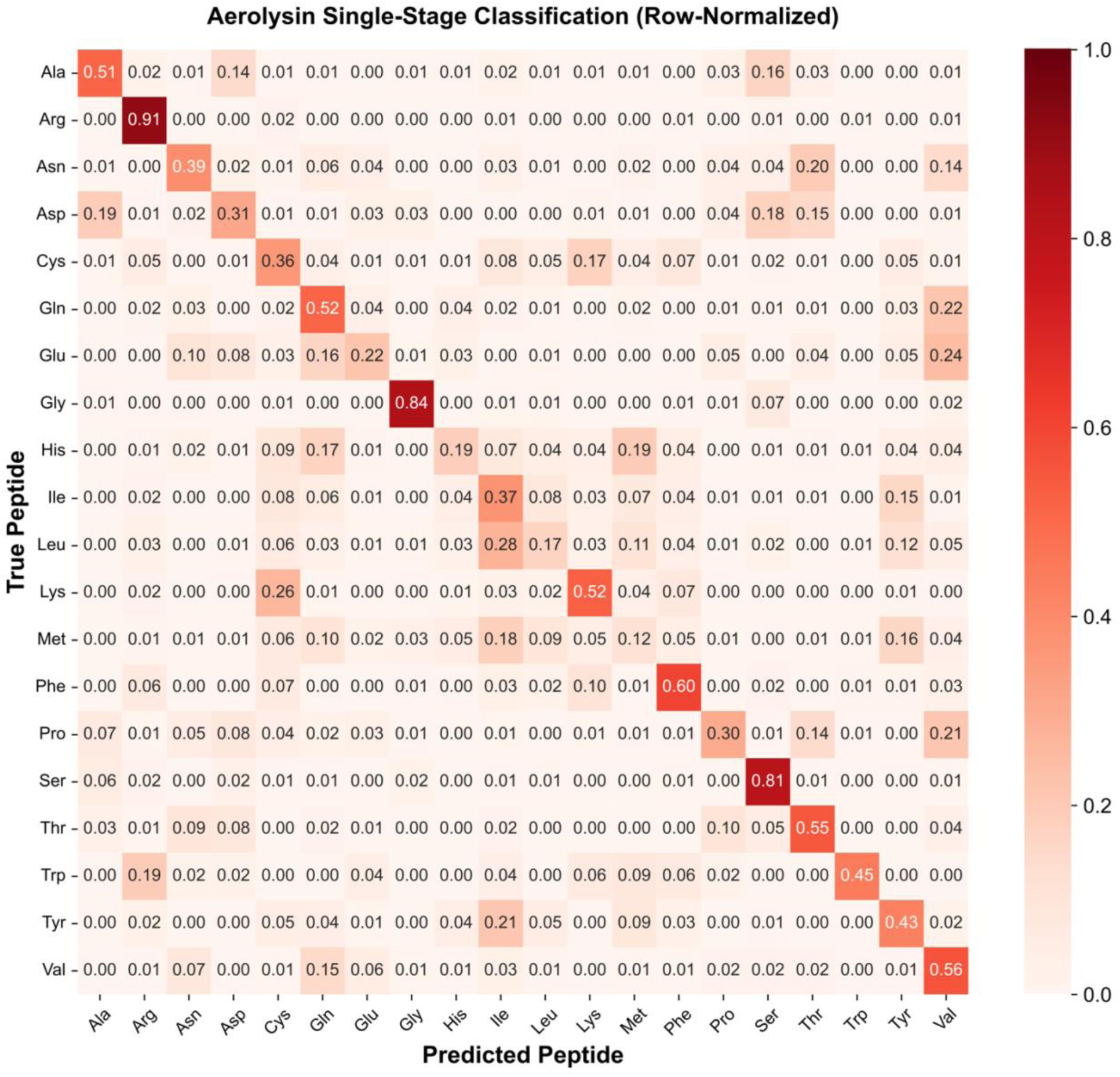
Machine Learning Benchmarking Exposes the Mathematical Ceiling of Static Nanopores. A 20-class, row-normalized confusion matrix evaluating the predictive accuracy of the static wild-type aerolysin nanopore (data derived from Ouldali et al., 2020) ^3^. Raw high-bandwidth (250 kHz) .abf recordings were processed using an automated 1D K-Means state-labeling algorithm and evaluated via a single-stage, multi-class XGBoost classifier. To eliminate human bias and expose the “Data Curation Fallacy,” no manual 2D thresholding filters were applied. Crucially, Cysteine evaluation strictly utilized recordings obtained in the presence of 25 mM DTT to ensure true monomeric translocation, preventing the artificial classification advantage of disulfide dimers in neutral pH conditions. To account for the extreme capture-rate disparities inherent to passive stochastic diffusion (e.g., Arginine events were significantly greater than Tryptophan events), the full, imbalanced dataset (>51,000 events) was utilized. Mathematically balanced sample weights were applied during model training to correct for this disparity without artificially starving the classifier via downsampling. Despite maximizing data density and applying algorithmic corrections, overall classification accuracy mathematically capped at 0.6692 (±0.0054) for three replicate train/test analyses. This empirical benchmark demonstrates that without the high-dimensional kinetic transition matrices natively generated by an active, dynamical translocase, static volumetric measurement is fundamentally insufficient to resolve isobaric and isosteric overlaps in a complex 20-class mixture, rendering it unviable for *de novo* sequencing.

## 7. Dismantling Notion that Sequence-to-Signal Cannot be Mapped by Machine Learning

A common computational critique of dynamical translocases is that their multistate gating behavior represents stochastic noise rather than sequence dependent signal, lacking the interpretability required for sequence-to-signal prediction. This view assumes that sequence-to-signal relationships can only be learned from simple low-dimensional observables. This perspective overlooks advances in computational biophysics, machine learning, and advanced deep neural networks, which have proven remarkably effective at extracting information from high-dimensional dynamical systems. The continuous fluctuations of the PA ϕ-clamp are not random noise; they are stochastic manifestations of and underlying energy landscape **(Fig. 2)**. By deploying a Physics-Informed Machine Learning (PIML) framework ^37,38^, we treat the translocase as a physically interpretable system rather than an opaque “black box.” The resulting models explicitly extract quantitative descriptors of the translocation event. A single peptide translocation yields a >60-dimensional feature vector, encompassing Markovian dwell time transition matrices, discrete state probabilities, dwell-time variances, conductance levels, and transition frequencies ^17,18^.

Far from being theoretically intractable, sequence-to-signal decoding is already an empirical reality. To directly compare dynamical and static sensing architectures, we deployed a single-stage baseline XGBoost classifier ^39^ on the full 20-canonical-amino-acid guest-host peptide (K_5_XXSXX) library (>550,000 events). To ensure a fair comparison, we intentionally matched the classifier architecture for aerolysin benchmarking, allowing differences in sensing performance to be attributed primarily to the underlying biological hardware. Tested under a 35 ms minimum-dwell time constraint to isolate deep, sustained multistate thermodynamic friction, the dynamic PA translocase achieved a robust 90.7% single-stage classification accuracy (**Fig. 5**). Crucially, the classifier separated isobaric residues, yielding F1-scores of 0.93 for leucine and 0.92 for isoleucine – targets that are notoriously difficult to distinguish using purely volumetric measurements and remain challenging for conventional proteomic workflows.

**Figure 5.**
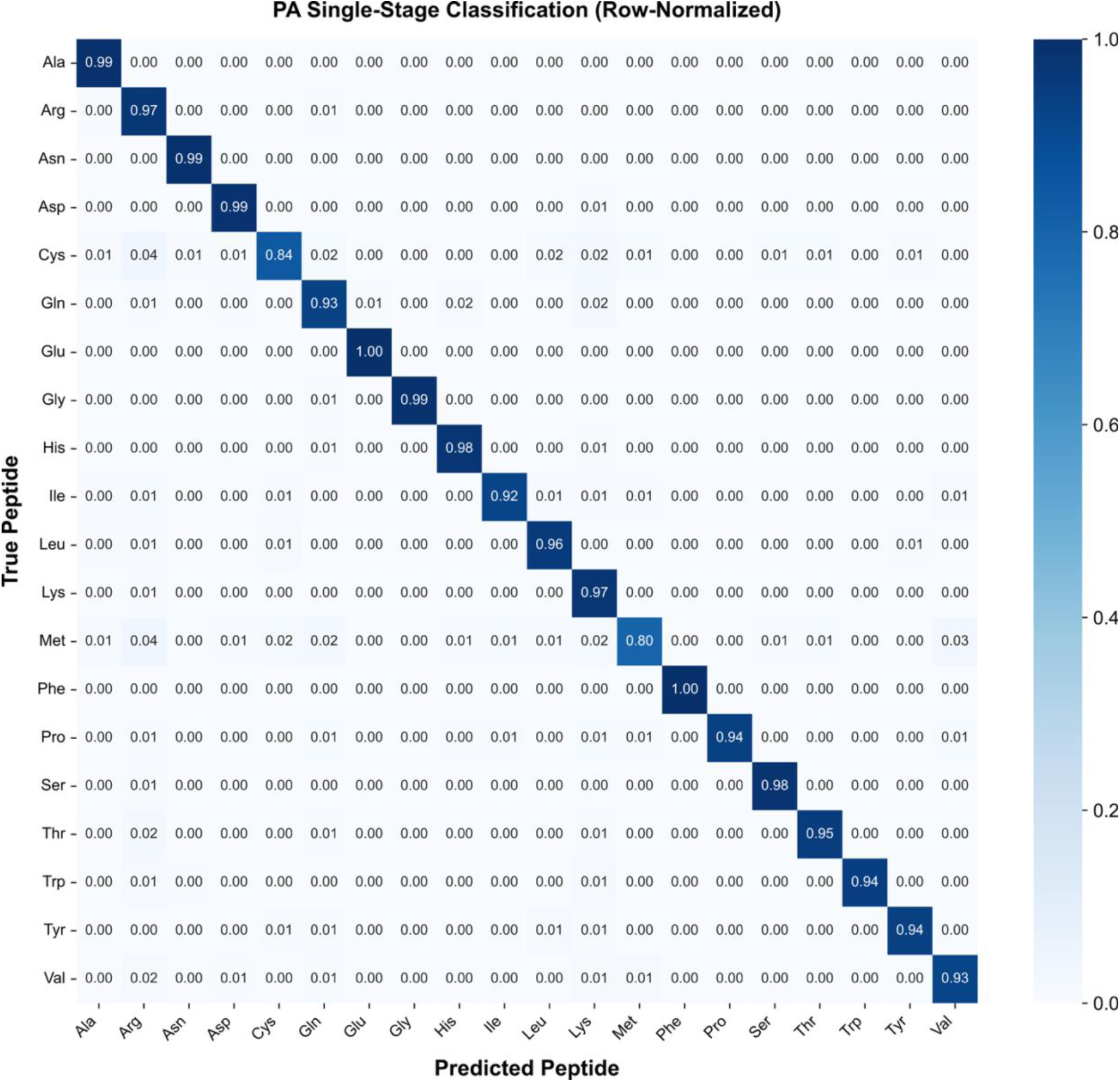
Resolving the Unresolvable via Dynamical Nanopore Hardware and PIML. A row normalized classification matrix generated via a single-stage baseline XGBoost classifier utilizing dynamic feature extraction from single-channel PA recordings. The dataset is entirely new data from at least three single-pore/membrane replicates derived from guest-host peptides compromising a full 20 amino acid class set and highlighting the platform’s capacity to discriminate between complex, highly similar peptide analogs. The host peptide sequence, concentrations, and nanopore conditions are given in Fig. 1C. Overall model accuracy is 0.9060 (±0.0155) for >550,000 total translocation events. Crucially, the PIML architecture effortlessly resolves isobaric residues (e.g., Leu vs. Ile) – targets possessing identical spatial volumes and masses that fundamentally confound conventional static nanopores and standard mass spectrometry.

The successful classification of a 20-amino-acid analyte library demonstrates that dynamical nanopore signals are amenable to sequence-to-signal modeling. Whereas static-pore projections require 200 ms peptide dwell times to approach 90% theoretical accuracy, the dynamical PA translocase achieves 90.7% empirical accuracy within 35 ms during continuous translocation. The static pore yields a low-dimensional representation of the analyte, whereas the dynamical translocase generates the high-dimensional kinetic information required for robust label-free protein discrimination.

## 8. Conclusion

Future progress in single-molecule proteomics will require sensing architectures designed for proteins rather than nucleic acids. Static nanopores have been extraordinarily successful for genomics, but when applied to chemically and conformationally diverse proteins they yield comparatively low-dimensional molecular fingerprints. In contrast, evolution has already produced a sophisticated protein-handling nanomachine in the anthrax toxin translocase. By recognizing the dynamic ϕ-clamp as a kinetic sensor and coupling it with Physics-Informed Machine Learning, translocation dynamics can be transformed into rich high-dimensional molecular fingerprints that extend beyond purely spatial measurements and chemical limitations of traditional platforms. The central challenge is no longer detecting proteins but extracting sufficient information to distinguish them.

Static pores function as fixed molecular calipers, extracting information primarily from excluded-volume measurements within a rigid constriction. Like any measuring device, a caliper can only report the information it is capable of sensing. As proteins increase in chemical and conformational complexity, these measurements yield only a limited set of geometric observables. Dynamical translocases expand this observable space through analyte-dependent conformational transitions, converting molecular interactions into high-dimensional kinetic fingerprints. The advantage is therefore not merely one of sensitivity, but of information content. Dynamical translocases generate far richer, analyte-dependent kinetic fingerprints that provide machine-learning algorithms with substantially more informative features for classification, discrimination, and ultimately sequence inference. The mathematical imperative for the field is clear: the future of proteomics lies in extracting richer information from dynamic molecular interactions.

## 9. Methods Summary

Single-channel electrophysiological recordings of guest-host peptide translocations through the anthrax toxin PA nanopore were acquired to map discrete, multi-state kinetic transitions. Translocation events were processed using a custom PIML framework, extracting a >60-dimensional vector of thermodynamic and temporal features per event (e.g., dwell time transition matrices, state probabilities, etc.) to train a multi-class XGBoost classifier ^39^ **(Table S1)**. To establish an objective, static-pore benchmark, legacy high-bandwidth aerolysin datasets ^3^ were re-analyzed using an unsupervised, uncurated machine learning pipeline to determine the empirical accuracy ceiling bounded by 1D excluded volume. Finally, an information-theoretic analysis (Shannon entropy) of mature translocase cargo (Lethal Factor and Edema Factor) was conducted against global proteomic baselines to mathematically map the evolutionary compositional bias required for active translocation.

Detailed protocols describing protein purification, electrophysiology, feature extraction matrices, ML hyperparameters, and bioinformatic scripts are provided in the Supplementary Information.

## Acknowledgments

The authors thank the department for stimulating discussions. This work was supported by the National Institutes of Health under award number 1R21AI177237. The content is solely the responsibility of the authors and does not necessarily represent the official views of the National Institutes of Health.

## Author Contributions

J.E.T. designed the experiments, collected data, analyzed results, and wrote the manuscript. P.S. contributed machine learning expertise and evaluated data analysis. B.A.K. conceived of the PIML pipeline, designed experiments, and wrote the manuscript.

## Code and Data Availability

The Ouldali et al. primary data utilized in this study were derived from publicly available datasets (DOI: 10.13012/B2IDB-4905767_V1). Anthrax toxin raw data, custom Python classes and scripts utilized for data processing, machine learning analysis, and figure generation are publicly available via Zenodo (DOI: 10.5281/zenodo.21453990). The basic preprocessing architecture and machine learning codebase are available on Github (https://github.com/bakrantz/Pept-Class).

## Declaration of AI and AI-Assisted Technologies in the Writing Process

During the preparation of this work, the author used Google Gemini and Google Search to assist with literature research, refine manuscript text for clarity and readability, and optimize custom computational code. After using these tools/services, the authors reviewed and edited the content as needed and takes full responsibility for the ultimate content and integrity of the publication.

