## Supplementary Information for "Beyond the Static Caliper: Dynamical Translocases and the Mathematical Imperative for Single-Molecule Proteomics"

**Running title:** Dynamical Nanopore Translocase Imperative

**Keywords:** Single-molecule proteomics, nanopore sequencing, Physics-Informed Machine Learning, translocase, anthrax toxin, Markov kinetics, aerolysin

### Materials and Methods

**Nanopore and peptides.** Monomeric 83-kDa PA ($PA_{83}$) preprotein was prepared as described [1]. $PA_{83}$ monomer was overexpressed in *Escherichia coli* BL21(DE3), using a pET22b plasmid, which directs expression to the periplasm. Cell cultures were grown at 37 °C in a custom 5 L fermentor using ECPM1 growth media [2], which was supplemented with carbenicillin (50 mg/L). Once reaching an $OD_{600}$ of 3-10, the cultures were then induced with 1 mM isopropyl β-d-thiogalactopyranoside for ~3 h at 30 °C. $PA_{83}$ was released from the periplasm by resuspending pelleted cells on ice using a wire whisk with 1 L of hypertonic sucrose buffer (20% sucrose, 20 mM Tris-Cl, 0.5 mM EDTA, pH 8) followed by osmotic shock of centrifuged/pelleted cells using a wire whisk in 1 L of hypotonic solution (5 mM $MgCl_2$). Released $PA_{83}$, isolated after centrifugation to remove cellular debris, was purified on Q-Sepharose anion-exchange chromatography in 20 mM Tris-Cl, pH 8.0 by binding and then eluting with a linear salt gradient using 20 mM Tris-Cl, pH 8.0 with 1 M NaCl.

To make nicked PA (nPA), purified $PA_{83}$ at a concentration of 1 mg/ml was treated with trypsin (1:1000 wt/wt trypsin:PA) for 30 min at room temperature. Trypsin was subsequently inhibited with soybean trypsin inhibitor at 1:100 dilution (wt/wt soybean trypsin inhibitor:PA). nPA was frozen in -80 °C in small aliquots to maintain reproducible nanopore insertion activity in planar bilayer experiments.

Ten-residue guest-host peptides of the general sequence, KKKKKXXSXX, where X = each of the 20 natural amino acids, were synthesized with standard L amino acids to >95% purity as indicated [3,4] (Elim Biopharmaceuticals). One stereochemical variant of X = W was produced (guest-host TrpDL), where instead of synthesizing the peptide with uniform L amino acids, an alternating pattern of D and L amino acids was used [4].

**Single-channel electrophysiology.** Planar lipid bilayer currents were recorded using an Axopatch 200B amplifier interfaced by a Digidata 1440A acquisition system (Molecular Devices) [1,4,5]. Membranes were formed by painting across a 150-μm aperture of a 1-mL polysulfone cup with 3% (wt/vol) 1,2-diphytanoyl-*sn*-glycero-3-phosphocholine (Avanti Polar Lipids) in *n*-decane. The *cis* (side to which the nPA is added) and *trans* chambers were bathed in symmetric universal bilayer buffer [6] with broad pH buffering range (UBB: 100 mM KCl, 1 mM EDTA, 10 mM oxalate, 10 mM MES, 10 mM phosphate, pH 5.60). Recordings were acquired at 600 Hz using PCLAMP10. The applied voltage is defined as $\Delta\psi = \psi_{cis} - \psi_{trans}$ (where $\psi_{trans}$ is defined as 0 mV).

Single-channel recordings of the guest-host peptide translocations via the PA nanopore were carried out as described [4] with some slight differences. A single PA channel was inserted into a painted bilayer at a Δψ of 70 mV by adding 10 ng of nPA (freshly diluted from a 1 mg/ml stock aliquot) to the *cis* side of the membrane. The activated nPA monomer dissociates into a 63-kDa portion that self-assembles into a heptamer prepore and converts to the nanopore state by inserting into the membrane in an oriented manner. Once a single channel inserted, the *cis* chamber was perfused by fresh UBB to remove excess uninserted nPA. Then the desired peptide analyte was added to the *cis* chamber at 20 nM. Translocation data were acquired at Δψ = +70 mV (cis positive), collecting recordings of the translocation event stream for up to one hour. At least three, replicate membranes with single pores were collected for each peptide analyte. Guest-host Cys peptide was also collected in isolation and in presence of equimolar amounts of TCEP as a control to test for disulfide bond intermediates that may have impeded translocation, albeit the working pH was probably too acidic at pH 5.6 for significant disulfide bond formation.

**Software and environment used for ML.** A Python environment was used with XGBoost (3.0.0) [7] and other standard modules. All Pept-Class source code is available at GitHub (https://github.com/bakrantz/Pept-Class).

**Data cleanup, downsampling and state labeling.** Minor processing as well as conductance state labeling of the raw single-channel event stream recordings was subsequently performed. Rare transient out-of-range current spikes, insertion of second channels, and inactivated channels were removed by a ‘force values’ routine in CLAMPFIT. Translocation recordings were acquired at 600 Hz but were downsampled to 400 Hz by decimation in Python using the scipy.signal library. 400 Hz was chosen to maximize the data volume at a consistent time step for ML/DL-based peptide classification to include earlier legacy data [8]. Four major discrete conductance states were detected in translocation event stream recordings using K-Means clustering, where baseline current drift was corrected by applying a moving window average offset (4000 time point window). While rare additional states were noted, the data were labeled for the four dominant states. By convention during event detection, the fully blocked peptide-bound state was state 0, the intermediate closest to the fully blocked state was state 1, the intermediate closest to the open state was state 2, and the open state was state 3. Start and stop times of all detected events were used to label the state of each time point in the raw current recordings, producing a three-column CSV file of the stream with columns labeled as ‘time’, ‘current’, and ‘state’. All labeled CSV stream files for the 20 natural amino acids as guests in the peptides were entered with all experimental metadata into a local annotated peptide database to aid in *in situ* loading/preprocessing for the tested ML model.

**Preprocessing, translocation event segmentation, and feature extraction.** Raw translocation event streams were preprocessed prior to segmentation and feature extraction. While the segmentation core can apply low- and high-pass filtering and baseline correction, these filters/corrections were not applied to the experimental guest-host peptide translocation datasets presented in this study. The most critical preprocessing parameter was the minimum event duration, which served as an effective filter for excluding very short-duration events.

Following the application of these processing parameters, the state-labeled raw event streams were segmented into individual translocation events. Each event was defined as initiating when the current changed from the fully open state (state 3, corresponding to baseline current) to any peptide-bound state (state 0, 1, or 2) and terminating when the current returned to the open state. From these segmented events, both raw current sequences and corresponding state sequences were extracted.

A comprehensive set of event-level physics informed features was then computed from these sequences using a custom segmentation core. This core maintains a generalizable framework to process peptide translocation events from systems exhibiting diverse mechanisms and an arbitrary number of states. These features were initially categorized into scalar, vector, and matrix data structures, with values in vector and matrix features being state or transition enumerated. Scalar features included: Shannon entropy of state sequence, event duration, number of transitions, time of the first transition, total number of states visited during the event, skewness and kurtosis. Vector features included: observed conductance state Boolean, observed conductance levels, probability of residing in each state, and longest dwell time in each state. Matrix features included: average dwell time for specific state-to-state transitions, variance of dwell time for transitions, and ratio of probabilities between states. The descriptions and dimensions of this feature set are in **Table S1**.

While the segmentation core also supports the extraction of global features that have yielded high-quality classification results in simulated datasets (e.g., >0.99 accuracy) [9], these were not utilized in the classifications of the experimental guest-host datasets presented here, as our focus was solely on the more challenging and practical task of individual event-level classification.

For downstream classification, all matrix features were flattened into one-dimensional arrays and appended with the vector and scalar features to form a single feature vector for each translocation event. These flattened descriptive key names for the features were generated to maintain traceability in subsequent applications. All processed event sequences, their flattened features, and associated feature key names were saved as a Python pickle object for efficient storage and retrieval. A local peptide events database was employed to track these preprocessed pickle files, thereby preventing redundant segmentations and feature extractions from raw datasets.

**Initial baseline single-stage ML guest-host peptide classification of anthrax toxin.** ML-based classification of peptide translocation events from single individual nanopore variants was performed using the gradient boosting framework, XGBoost [7], which was implemented closely to as described previously [8,10]. For this, peptide event data, previously extracted and characterized into event-level features, were loaded from a local database. The comprehensive dataset was then split into an 80% training set and a 20% testing set for model development and evaluation, respectively. The XGBoost classifier was configured with the following key parameters: a multiclass classification objective, the number of target classes set to the total number of peptides, 1000 boosting rounds (trees), and a learning rate of 0.05 to control the step size shrinkage. The model's performance during training was monitored using the multiclass classification error. The trained model's performance was evaluated on the unseen testing dataset. Classification metrics including accuracy, precision, recall, and F1-score were summarized in a standard classification report, and a confusion matrix was generated to visualize per-class prediction accuracy.

**ML re-analysis of static aerolysin nanopore datasets.** To establish an empirical benchmark against static nanopore platforms, legacy aerolysin datasets [11] encompassing 20 canonical amino acid target classes were re-analyzed utilizing an unsupervised machine learning pipeline. The peptide sequence was XRRRRRRR, where X was one of the 20 natural amino acids. The X=C peptide had several available datasets, but the DTT treated and, therefore, monomeric peptide set was used. Raw native binary .abf recordings originally sampled at 250 kHz were voltage-filtered, aggressively trimmed of initial capacitive transients (first 100 ms of each voltage-stepped sweep), and decimated to a 10 kHz bandwidth. Continuous pseudo-streams were generated utilizing 1D K-Means clustering to assign binary state labels (Open vs. Blocked). To preserve the true biophysical reality of the sensor and eliminate human bias in event selection, every contiguous blocked event was segmented without imposing minimum dwell time or amplitude thresholding. Because static pores strictly lack the dynamic active-site structural transitions observed in translocases, feature extraction was intrinsically limited to a basic set of scalar metrics: dwell time, mean current, standard deviation of current, variance, skewness, and kurtosis. These feature-starved event dictionaries were packaged into Python pickle objects for downstream processing. The extracted events were evaluated utilizing a multi-class XGBoost classifier. Due to the reliance on passive stochastic diffusion for capture, the resulting dataset exhibited uneven event counts per class. To evaluate the data rigorously without discarding tens of thousands of valid observations via downsampling, the full imbalanced dataset (>50,000 events) was utilized. Mathematically balanced sample weights were computed natively via sklearn and applied during XGBoost training to correct for capture-rate disparities.

**Evolutionary Bioinformatics and Information Theory.** To evaluate the evolutionary and compositional bias of anthrax toxin translocase cargo, the primary FASTA sequences for *Bacillus anthracis* PA, Lethal Factor (LF), and Edema Factor (EF) were analyzed. To ensure the analysis exclusively reflected the thermodynamic constraints of active translocation rather than bulk translation, N-terminal secretion signal sequences were surgically trimmed to yield the mature polypeptides (PA initiated at EVKQE; LF at AGGHG; EF at MNEHY). Baseline global proteomic amino acid frequencies were derived from the canonical McCaldon and Argos (1988) distribution [12]. Fractional occurrences ($p_i$) for all 20 standard amino acids were calculated for each mature toxin. Shannon Information Entropy (H) was computed for each sequence utilizing the equation $H = -\Sigma\, p_i \log_2(p_i)$. The specific evolutionary fingerprint of the cargo was visualized by calculating the $Log_2$ fold-change of each amino acid's frequency relative to the global proteome baseline. All bioinformatic sweeps and visualizations were executed using standard Python libraries (pandas, numpy, matplotlib).

**Table S1. Physics-Informed Feature Engineering Definitions[1].**

| Feature Category | Specific Feature Name | Data Structure | Feature Count (N=4) | Biophysical Description |
|---|---|---|---|---|
| Scalars | Event Duration | Scalar | 1 | Total time from capture (entry) to release (exit). |
| | Number of Transitions | Scalar | 1 | Total count of state switches (measure of flickering). |
| | Time to First Transition | Scalar | 1 | Duration of the initial state entry (often State 1 or 0). |
| | Total States Visited | Scalar | 1 | Measure of how much of the energy landscape was explored (1–4). |
| | Shannon Entropy | Scalar | 1 | Information content of the state sequence (complexity metric). |
| | Skewness | Scalar | 1 | Asymmetry of state occupancy (measure of energetic bias favoring deeper or shallower blockades). |
| | Kurtosis | Scalar | 1 | Propensity for extreme transient excursions (measure of rare, heavy-tailed flickering events). |
| State Vector[2] | Observed State Boolean | $1\times N$ Vector | 4 | Binary flag (0/1) indicating if State i was visited. |
| | Observed Conductance | $1\times N$ Vector | 4 | Mean current level for State $i$ using scaled current. |
| | State Probability | $1\times N$ Vector | 4 | Fractional occupancy (time spent in State $i$ / total duration). |
| | Longest Dwell Time | $1\times N$ Vector | 4 | The maximum single dwell duration observed for State $i$. |
| Transition Matrix[2] | Mean Transition Dwell | $N\times N$ Matrix | 16 | Average time spent in State $i$ before transitioning to State $j$. |
| | Dwell Time Variance | $N\times N$ Matrix | 16 | Variance of dwells for specific $i\rightarrow j$ transitions. |
| | Probability Ratios | $N\times N$ Matrix | 16 | Ratio of occupancy probabilities ($P_i/P_j$) between all state pairs. |
| **Total Features** | | | **71** | |

[1]A total of 71 biophysical features were extracted for each translocation event based on the 4-state kinetic model (States 0, 1, 2, 3) observed in the PA nanopore.

[2]Features corresponding to states or transitions not observed in a specific event (e.g., if State 0 is never visited, or a specific 0→2 transition never occurs) are assigned NaN (not a number). The XGBoost algorithm natively handles these missing values, utilizing them as informative signals regarding the unvisited regions of the energy landscape.
